# Sequential genetic and epigenetic alterations in human pluripotent stem cells for recurrent abnormality

**DOI:** 10.1101/2023.12.17.572095

**Authors:** Yun-Jeong Kim, Byunghee Kang, Solbi Kweon, Sejin Oh, Dayeon Kim, Dayeon Gil, Hyeonji Lee, Jung-Hyun Kim, Ji Hyeon Ju, Tae-Young Roh, Chang Pyo Hong, Hyuk-Jin Cha

## Abstract

Human embryonic stem cells (hESCs) are naturally equipped to maintain genome integrity to minimize genetic mutations during early embryo development. However, genetic aberration risks and subsequent cellular changes in hESCs during *in vitro* culture pose a significant threat to stem cell therapy. While a few studies have reported specific somatic mutations and copy number variations (CNVs), the molecular mechanisms underlying ‘culture-adapted phenotype’ acquisitions of hESCs are largely unknown. Therefore, we conducted comprehensive genomic, single-cell transcriptomic, and single-cell ATAC-seq analyses of an isogenic hESC model displaying definitive ‘culture-adapted phenotypes.’ Notably, hPSCs with a copy number gain of 20q11.21 during early passage did not present ‘culture-adapted phenotypes’ nor *BCL2L1* induction. Single-cell RNAseq and ATACseq analyses revealed active transcriptional regulation at 20q11.21 loci at late-passaged hESCs with the induced *BCL2L1* and *TPX2* to trigger ‘culture-adapted phenotypes’ was associated with epigenetic changes facilitating TEA domain (TEAD) binding. These results suggest that copy number 20q11.21 gain and additional epigenetic changes are necessary for expressing ‘culture-adapted phenotypes’ by activating gene transcription at this specific locus.

## Introduction

Human pluripotent stem cells (hPSCs) are a primary cell source for stem cell-based regenerative therapy due to their unique pluripotency. Owing to extensive preclinical research, many clinical studies for Parkinson’s disease, macular degeneration, heart failure, and some cancers are underway (Deinsberger *et al*, 2020). However, other than potential teratoma formation by unintended residual undifferentiated hPSCs (Jeong *et al*, 2017), unexpected genetic aberrations in hPSCs (Mandai *et al*, 2017) are predominant technical hurdles for ensuring safe stem cell therapy due to their clinical uncertainty (Andrews *et al*, 2017).

Considerable attention has been garnered for understanding the biological consequences, risk thresholds, and causes of hPSC genetic instability, as numerous studies report recurrent copy number variations (CNVs) in various chromosomes, including 20q11.21 (Andrews *et al*., 2017). Genetic hPSC aberrations vary from CNVs, somatic mutations, and chromosome alterations (e.g., trisomy at the 12 and 17 chromosomes) (Ben-David *et al*, 2014; Lund *et al*, 2012). Nevertheless, somatic hPSC mutations during *in vitro* culture are relatively rare (Kuijk *et al*, 2020; Thompson *et al*, 2020). Thus, the collapse of unique hPSC characteristics, especially their high susceptibility to genotoxic stimuli (Dumitru *et al*, 2012; Liu *et al*, 2013) to preserve genome integrity, is closely associated with genetic aberration. This observation may explain the frequent 20q11.21 loci gains containing *BCL2L1* (International Stem Cell *et al*, 2011) to induce anti-apoptotic BCL-xL protein (Avery *et al*, 2013; Cho *et al*, 2018; Nguyen *et al*, 2014) and 17p13.1 loss where TP53 is present (Amir *et al*, 2017) and somatic TP53 mutations (Merkle *et al*, 2017) in hPSCs during prolonged *in vitro* culture. Interestingly, these genetic aberrations such as the induction of BCL-xL as an ‘acquired survival trait’, serve to rescue cells from apoptosis triggered by aberrant mitosis, often resulting from TPX2 induction in hPSCs (Jeong *et al*, 2023). This, in turn, lead to aneuploidy or additional aberrations (Zhang *et al*, 2019). Consequently, genetic alterations that enable survival under the selective pressures of *in vitro* culture conditions can potentially trigger further structural changes in the genome (Halliwell *et al*, 2020). The hypothesis has been substantiated through retrospective studies with multiple hPSC lines collected from diverse research institutions (International Stem Cell *et al*., 2011; Merkle *et al*., 2017).

However, the 20q11.21 gain alone is insufficient to drive precise phenotypic changes such as ‘acquired survival advantage (Jo *et al*, 2020).’ Thus, additional cue would be necessary to induce *BCL2L1* and *TPX2* gene transcription, which yields clear ‘acquired survival traits’ through BCL-xL protein and YAP/TEAD4-dependent gene expressions, respectively (Avery *et al*., 2013; Cho *et al*., 2018; Kim *et al*, 2023; Nguyen *et al*., 2014). Recurrent epigenetic changes, such as hyper-/hypo-methylation, parental imprinting loss, and variable X chromosome inactivation, transpire during prolonged hPSC cultures (Bar & Benvenisty, 2019). In particular, hypermethylation begets the epigenetic repression of multiple antioxidant genes (Konki *et al*, 2016) in prolonged hPSC cultures with abnormal karyotypes, ‘differentiation-related genes,’ and ‘tumor-suppressor genes’ (Weissbein *et al*, 2017). Therefore, isogenic hESC sets with varying consecutive durations during *in vitro* maintenance would exhibit unique cellular and molecular characteristics, enhancing progressive deterioration monitoring.

This study used an isogenic human embryonic stem cell (hESC) set with different culture periods of up to six years to monitor stepwise epigenetic and genetic alterations during consecutive *in vitro* hESC cultures. Multi-omics analysis, including whole genome sequencing (WGS), single-cell RNA sequencing (scRNA-seq), and single-cell ATAC sequencing (scATAC-seq), revealed that dominant-negative TP53 mutations accentuated somatic mutations. In addition, epigenetic alterations at 20q11.21 loci promoted gene expression at 20q11.21 through transcriptional enhanced associate domain (TEAD)-dependent transcription. These results account for 20q11.21 gain and TP53 mutations, the most prevalent genetic aberrations in hPSCs.

## Materials and Methods

Detains of the methods are available in the online supplement.

### Cell line and culture

Human embryonic stem cell (WA09: H9, WiCell Research Institute) were maintained in iPSC-brew StemMACS (Miltenyi biotechnology, #130-104-368) with 0.1% gentamycin (Gibco, Waltham, MA, USA, #15750-060) on a matrigel (Corning, Corning, NY, USA, #354277)-coated cell culture dish at 37°C and humidified to 5% in a CO2 incubator. Cells were maintained with daily changed media and passaged every 5-6 days. Upon transfer, hESCs were rinsed with DMEM/F-12(Gibco) and detached enzymatically with Dispase (Life Technologies) followed by 3 times washed with DMEM/F-12 (Gibco #11320-033) media before plating. If needed, 10 µM of Y27632 (Peprotech#1293823) was used for cellular attachment.

### Whole-genome sequencing and detection of somatic mutations

Genomic DNA was isolated from P1, P2, P3, and P4 hESCs using the Wizard® HMW DNA Extraction Kit (#A2920, Promega, USA) for whole-genome sequencing on the Illumina NovaSeq platform. Libraries were prepared from 1 µg of input DNA using a TruSeq DNA Sample Prep Kit according to the manufacturer’s instructions (Illumina, Inc., San Diego, CA, USA). The sheared DNA fragments underwent end-repair, A-tailing, adaptor ligation, and amplification, followed by clean-up. These libraries were then subjected to paired-end sequencing with a read length of 150 bp on the Illumina NovaSeq 6000 platform, yielding an average of 106.1 Gb per library.

Clean reads from each sample, exhibiting a quality of >Q30 (%), were aligned to the human reference genome (GRCh37/hg19) utilizing BWA (version 0.7.17). PCR duplicates were identified and removed via MarkDuplicates of the Picard tool (version 2.25.5). To optimize read mapping quality, base quality score recalibration was conducted using the BaseRecalibrator Tool in GATK (version 4.1.9.0). Somatic mutations in P2, P3, and P4 hESCs were detected using MuTect2 (version 4.1.8.1), with P1 hESC serving as the reference. High-confidence somatic mutations were characterized as the subset of somatic variant calls fulfilling the following criteria: (1) sites with a VAF of 0 in P1 hESC, (2) biallelic SNVs, (3) mapped read depth of ≥ 20, (4) a minimum alternate read depth of ≥ 3, (5) exclusion of germline-like heterozygous SNVs, and (6) exclusion of SNVs on the X and Y chromosomes. The identified somatic variants were annotated with SnpEff (version 4.3) and cross-referenced against the Catalogue Of Somatic Mutations In Cancer (COSMIC) and The Cancer Genome Atlas (TCGA) databases.

### Evaluating tumorigenic potential in hESC with missense mutations in TP53-Linked genes

To assess the extent of missense mutations occurring in TP53-linked genes, we developed a scoring system based on two hypotheses. First, we assumed that the impact of a mutation is related to inter-amino acid change frequency. For instance, the KRAS-G12D mutation, known to cause tumors, involves a change from neutral glycine to negative aspartic acid. According to the BLOSUM100 matrix, this mutation occurs naturally with approximately 1.8% probability. Therefore, we derived scores for amino acid changes based on the BLOSUM100 matrix for mutations occurring in sequences with 100% similarity. We summed the scores for amino acid changes with negative values, assuming that changes occurring frequently would not affect the gene’s function, to create the BLOSUM score (B-score) (Equation 1).

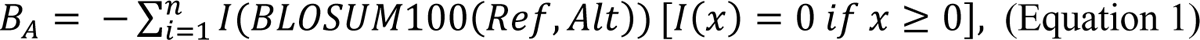

where *B*_*A*_ is B-score of sample A. *BLOSUM100*(*Ref*, *Alt*) represents BLOSUM100 score for amino acid transition between reference amino acid (Ref) and altered amino acid (Alt). *I(x)* is an indicator function that returns 0 when there are positive values in the inter-amino acid change according to the BLOSUM100 matrix.

On the other hand, there are mutations like TP53-R135H, a well-known tumorigenic TP53 hotspot mutation, that do not follow the first hypothesis, where both R and H are neutrally charged and occur with about 27% probability. To complement this, the second hypothesis introduced the possibility of the mutation being a hotspot mutation. For this scoring system, we obtained *p*-values from a study by Gao et al. (2017). We transformed these *p*-values into log_2_-scale and, if the mutation matched Gao’s defined list, summed the log_2_(*p*-value) values to create the Hotspot score (H-score) (Equation 2).

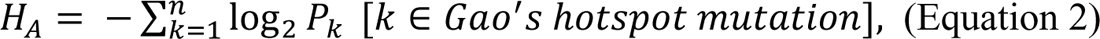

where *H*_*A*_ is H-score of sample A and *P*_*k*_ represents *p*-values of genes that have Gao’s hotspot mutations.

Finally, the *TP53score*_*A*_ for sample A’s TP53-linked gene was calculated by summing the B-score and H-score, and then passing the value through the hyperbolic tangent function to produce a standardized score ranging from 0 to 1 (Equation 3).

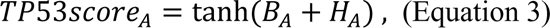

### Copy number variation (CNV) detection in whole-genome sequence data

CNVs were identified in the whole-genome sequence data of P2, P3, and P4 hESCs using CNVKit (version 0.9.10), with P1 hESC serving as the reference. CNVs exhibiting significant gains or losses were selected, applying a stringent cutoff of an absolute copy number ratio exceeding 0.85. Furthermore, CNVs identified in the whole-genome sequence data were confirmed using the CytoScan HD array (Thermo Fisher Scientific) or array comparative genomic hybridization (array CGH) (Thermo Fisher Scientific).

### Bulk RNA-seq and bioinformatic analysis

cDNA libraries were prepared from 1 μg total RNA of each sample using the TruSeq Stranded mRNA Sample Prep Kit (Illumina, Inc., San Diego, CA, USA), according to the manufacturer’s instructions. After qPCR validation, libraries were subjected to paired-end sequencing with a 150 bp read length using an Illumina NovaSeq 6000 platform, yielding an average of 5.9 Gb per library.

Clean reads with quality scores of >Q30 were aligned to the human reference genome (GRCh37/hg19) using STAR (version 2.7.1a) with the default parameters. Gene expression quantitation was performed using RSEM (version 1.3.1). Transcripts Per Kilobase Million (TPM) was calculated as the expression value. The differentially expressed genes (DEGs) were identified by DESeq2 (version 1.26.0) with the cutoff set at *q* < 0.05 and > 1.5-fold change. Gene ontology (GO) enrichment analysis for DEGs was conducted using the DAVID Gene Functional Classification Tool (http://david.abcc.ncifcrf.gov; version 6.8) with the cutoff of an EASE score < 0.01.

### Single-cell RNA sequencing (scRNA-Seq) and bioinformatic analysis

Single-cell suspensions with 90% cell viability were processed on the 10x Chromium Controller using Chromium Next GEM Single Cell 3’ Reagent Kits v3.1 (10x Genomics) according to the manufacturer’s instructions. Cells were partitioned into nanoliter-scale Gel Beads-in-Emulsion (GEMs) with target recovery of 10,000 cells. The single-cell 3 prime mRNA seq library was generated by reverse transcription, cDNA amplification, fragmentation, and ligation with adaptors followed by sample index PCR. Resulting libraries were quality checked by Bioanalyzer and sequenced on an Illumina NovaSeq 6000 (index = 8 bases, read 1 = 26 bases, and read 2 = 91 bases). We generally acquired an average of 516.1 million reads per library.

Paired-end sequencing reads were processed by Cell Ranger (10x Genomics software, version 1.3.1). Reads were aligned to the human reference genome (GRCh37/hg19) for demultiplexing, barcode filtering, gene quantification, and gene annotation. Barcodes with less than 10% of the 99th percentile of total unique molecular identifier (UMI) counts per barcode, which are likely to be empty droplets, were removed. Following this quality control step, gene barcode matrices for each sample were generated by counting the number of UMIs for a given gene in the individual cell.

To analyze heterogeneous composition of cell populations, gene barcode matrices were processed with Seurat R package (version 2.4). Low-quality cells with less than 1000 or more than 7200 detected genes were removed. In addition, cells with their mitochondrial gene content of >11% were also removed. After filtering out low-quality cells, the gene expression value was then natural log-transformed and normalized to sequencing depth scaled by a multiplier (e.g., 10,000). To reduce the variance caused by unwanted sources, variations in gene expression driven by cell cycle stages (S and G2/M phase) and mitochondrial gene expression were regressed out using the vars.to.regress argument in the ScaleData function of Seurat. Next, unsupervised clustering analysis for scRNA-Seq dataset was performed. Highly variable genes were selected using the FindVariableFeatures function in Seurat, and dimensionality reduction on the data was performed by computing the significant principal components on highly variable genes. Unsupervised clustering was performed by using the FindClusters function in Seurat with the resolution argument set to 0.3, and the resulting clusters were then visualized in a Uniform Manifold Approximation and Projection (UMAP) plot. After unsupervised clustering, DEGs among each cell cluster were identified using the

FindAllMarkers function in Seurat. Additionally, significant DEGs were filtered based on the following criteria: (1) *p* < 0.001, (2) a fold change of greater than an absolute 0.584 in log2-scale, and (3) differences in expression between pct1.1 and pct.2 exceeding 0.3. To annotate the biological process functions of DEGs from clusters, GO enrichment analysis was conducted using the DAVID Gene Functional Classification Tool (http://david.abcc.ncifcrf.gov; version 6.8), applying a cutoff of an EASE score < 0.01.

We also used the Monocle3 R package to reconstruct cellular trajectories by computing and ordering the sequence of gene expression changes of the cells collected from different time points (passaged *in vitro* <50, around 100 times, 200s, and 300s hESCs). First, DEGs were identified using differentialGeneTest function with *q* < 0.01 in Monocle3. The dimensions of the data were reduced using the Discriminative Dimensionality Reduction with Trees (DDRTree) method. Next, the cells based on the selected DEGs were ordered using orderCells function in Monocle3 and the trajectory of the cells was visualized by the plot_cell_trajectory function in Monocle3.

### Single-cell ATAC-seq library preparation and bioinformatic analysis

Nuclei were isolated from cells using the Nuclei Isolation Kit: Nuclei EZ Prep (NUC101-1KT) according to the manufacturer’s instructions. Sequencing library was prepared using Chromium Next GEM Single Cell ATAC Reagent Kit (10x Genomics; PN-1000176, v1.1). About 8,000 nuclei were isolated for Tn5 transposition, nuclei barcoding, and library construction. Quality checked libraries were sequenced using the Illumina Sequencing platform. The paired sequencing reads and barcodes were demultiplexed, preprocessed, aligned to hg38 human reference genome and used to call peaks with single-cell ATAC Cell Ranger pipeline (version 2.0.0). Resulting matrices were further analyzed using R package Signac (version 1.4.0) (Stuart *et al*, 2021). Clustering cells and finding differentially accessible regions (DARs) were performed with integrated samples. DARs were selected with following threshold: false discovery rat < 0.01 and log_2_ fold change > 0.3. All collected DARs among major clusters were grouped by K-means clustering with k=10. Cis-coaccessibility network (CCAN) was built using Cicero (version 1.3.4.11) with co-accessibility score > 0.05 as threshold (Pliner *et al*, 2018). Genes with upstream 1kb region included in CCAN were collected as candidate targets for DARs. Functional enrichment assay of candidate target genes was performed using metascape (Zhou *et al*, 2019). TF activity score was calculated by RunChromVAR() function (Schep *et al*, 2017) with JASPAR2020 database (Fornes *et al*, 2020). The AUC score was calculated by FindMarkers() function. To profile each cell clusters, subset-bam (version 1.1.0) was used to separate read pairs from integrated sequencing data. Peak calling for each cluster was performed by MACS2 (version 2.2.9.1) (Zhang *et al*, 2008). ATAC-seq footprints were analyzed within peaks using HINT-ATAC (version 1.0.2) (Li *et al*, 2019). The deepTools (version 3.1.3) (Ramirez *et al*, 2016) was used to calculate fold change of ATAC-seq signal against P1 sample and to draw heatmap around peak centers. To compare ATAC and CNV, the patterns of changes in ATAC and CNV were calculated across each gene body using log_2_ fold change against P1. The similarity scores of ATAC and RNA were calculated by inverse of Euclidean distance between pattern of ATAC and RNA.

## Results

### The drastic somatic mutation increase in culture-adapted hESCs

We have identified the cellular and molecular events during long-term *in vitro* passage using the H9 hESC set with different passage numbers, maintained up to 6 years (Fig. 1A). Genomic variants (i.e., P3 and P4 hESCs) share typical culture-adapted phenotypes, as previously demonstrated (Avery *et al*., 2013; Markouli *et al*, 2019; Nguyen *et al*., 2014; Price *et al*, 2021; Spits *et al*, 2008; Zhang *et al*., 2019) and depicted in Figure 1A. As these culture-adapted phenotypes were manifested in P3 hESCs (but not P2 hESCs), P4 hESCs with an additional 17q24 gain (Jeong *et al*., 2023) (Fig. S1A) from P3 hESCs (with a 20q11.21 gain) encourages closer examination of ‘stepwise variation models for genome hESCs instability (Ben-David, 2015; Halliwell *et al*., 2020).’

**Figure 1.**
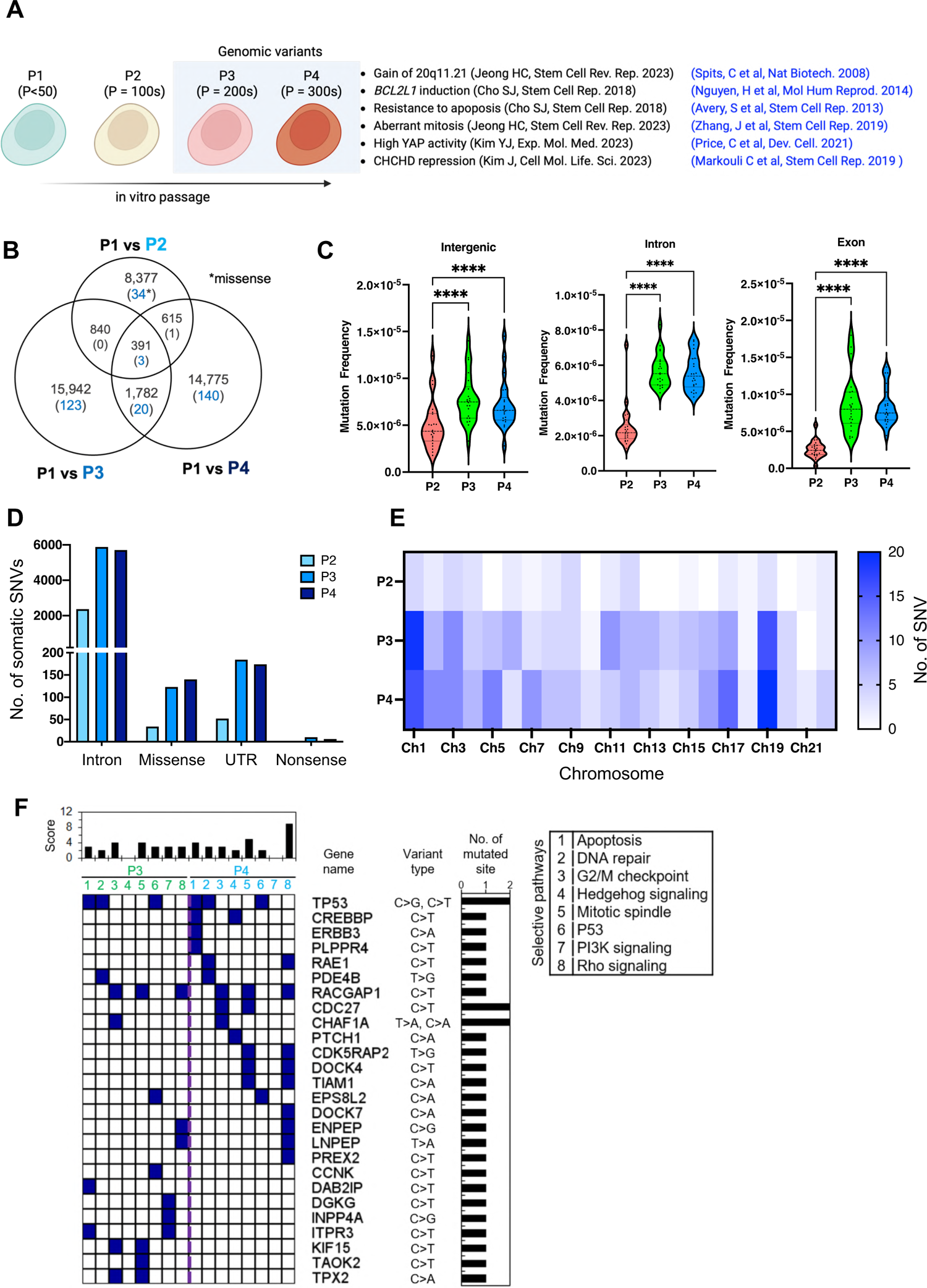
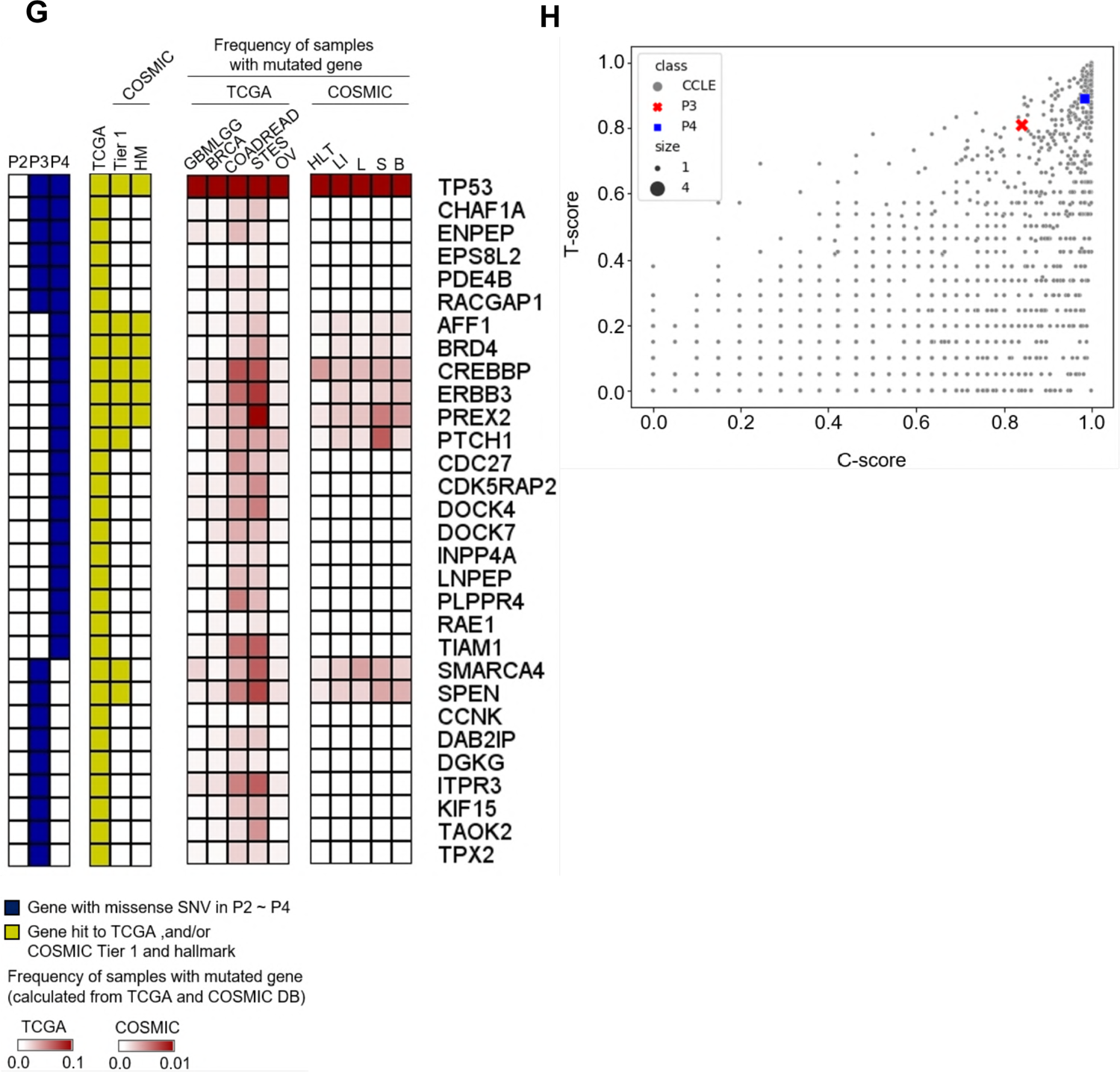
The drastic somatic mutation increase in culture-adapted hESCs (A) General scheme of passage dependent isogenic pair of hESCs. P1 hESCs: lesser 50 passages (one year culture), P2 hESCs: over 100 passages (two years culture), P3 hESCs: over 200 passages (four years culture), and P4 hESCs: over 300 passages (over six years culture). (B) Number of somatic single nucleotide variants (SNVs) detected by whole-genome sequencing of P1, P2, P3 and P4 hESCs, and overlaps of somatic SNVs(Detected using Mutect2) among those hESCs. The number of missense SNVs was highlighted in blue. (C) The frequency of somatic SNVs in intergenic regions, introns, and exons in P2, P3 and P4 hESCs were calculated by dividing the number of somatic SNVs found in each genomic feature by the length of that region. (D) The number of somatic intron, missense, 5’/3’-UTR, and nonsense SNVs identified in P2, P3, and P4 hESCs. (E) The chromosomal distribution of somatic SNVs in P2, P3, and P4 hESCs. The abundance of somatic SNVs was presented with a heatmap. (F) Functional pathways involved in the functions of survival and proliferation in stem cells, and mutated in P3 and P4 were analyzed using QIAGEN Ingenuity Pathway Analysis as described in the supplementary method. (G) Genes with somatic missense SNVs (highlighted in blue in the first panel of Figure 1G) were checked against The Cancer Genome Atlas (TCGA) and the Catalogue of Somatic Mutations in Cancer (COSMIC)-Tier 1 and -hallmark (HM) databases (highlighted in yellow in the second panel of Figure 1G). (H) Somatic missense SNVs in P3 and P4 hESCs were searched against the TP53-directed network and/or COSMIC Tier 1 datasets, and the impact of those mutations was assessed using T-scores for TP53-direct network and C-scores for COSMIC Tier 1 (see the Methods). Finally, the resulting scores were compared to data from the Cancer Cell Line Encyclopedia (CCLE).

We investigated progressive somatic mutations in long-term culture-adapted hESCs by performing whole genome sequencing (WGS) on the hESCs set and analyzing the genomic distribution of somatic single-nucleotide variants (SNVs) in P2, P3, and P4 hESCs with P1 as a normal control. Following quality control guidelines (see the Materials and Methods section), 42,722 somatic SNVs were identified. The primary variant types analyzed were C>T, C>A, and T>C (Fig. S1B). Somatic SNVs in P3 and P4 hESCs increased by approximately 1.8-fold compared to P2 hESCs, coinciding with the significant occurrence of P3 and P4-specific SNVs. (Fig. 1B). This correlation demonstrates an association between the increase in passage number and somatic mutation.

Intergenic, intronic, and exonic region mutation frequencies, as annotated in the GRCh37 genome assembly, were measured and normalized to base-pair content of each feature class (Fig. 1C, S1B, and S1C). P2 and P3/P4 hESC mutation frequencies significantly differed in introns (approximately 2.53 [*P*= 2.25 x 10^-8^] and 2.47 [*P*= 3.26 x 10^-9^] times more abundant in P3 and P4 than in P2, respectively) and exons (approximately 4.10 [*P*= 4.58 x 10^-8^] and 4.36 [*P*= 1.07 x 10^-8^] times more abundant in P3 and P4 than in P2, respectively). Specifically, concerning coding regions, a significant increase in missense and nonsense variants was observed in P3 and P4 (Fig. 1D). Our results indicate that genetic variant accumulation in genic regions during long-term in vitro passage augments hESC genome instability.

Interestingly, we observed that somatic mutations were non-random; somatic SNVs in chromosomes 1 and 19 were the most prevalent (Fig. 1E and S1D). Notably, recurrent gains were frequently observed in chromosome 1 (Narva *et al*, 2010) (Martins-Taylor *et al*, 2011). Along with the drastic increase in P3 and P4 hESC mutations, we identified 29 missense mutations, primarily consisting of C>T and C>A variant types (Fig. S1C), in 26 genes associated with apoptosis, DNA repair, G2/M checkpoint (cell cycle), mitotic spindle, P53, Hedgehog signaling, PI3K signaling, and Rho signaling (Fig. 1F). Among these pathways, PI3K gain and Rho signaling loss also engender the survival benefit of hPSCs from unbiased genome-wide screening (Weissbein *et al*, 2019).

We identified 30 genes with missense SNVs in P3 or P4 hESCs associated with tumorigenesis in The Cancer Genome Atlas (TCGA) (second panel in Fig. 1G). These genes were frequently localized in colon and rectum adenocarcinoma in TCGA (third panel in Fig. 1G). Additionally, nine of these genes, including *TP53*, *AFF1*, *BRD4*, *CREBBP*, *ERBB3*, *PREX2*, and *PTCH1*, were also prevalent in the Catalogue of Somatic Mutations in Cancer (COSMIC) (Tate *et al*, 2019) Tier 1. COSMIC collects genes contributing to tumor promotion and directly accounts for clinical PSC cell therapy risks (second and fourth panels in Fig. 1G). To further assess tumorigenesis potential in P3 and P4 hESCs, we scored the impact of missense SNVs in genes within the TP53 network (T-score) or COSMIC Tier 1 (C-score) based on the Cancer Cell Line Encyclopedia (CCLE; Fig. 1H). The assessment revealed high tumorigenesis potential in P3 and P4 hESCs, as indicated by their high T- and C-scores, where the closer the score is to 1, the higher the impact it represents (Fig. 1H).

### TP53 mutations relative to an increase in somatic mutations

Recent studies have highlighted the incidence of dominant-negative mutations in TP53 within hPSCs (Merkle *et al*., 2017). Since p53 is pivotal in inducing cell death in hPSCs (Lee *et al*, 2013; Liu *et al*., 2013), p53 stabilization markedly begets this outcome. Thus, with passaging, clonal dominancy is readily achieved in TP53 mutant clones (Amir *et al*., 2017). Two somatic missense SNVs, including c.524G>A (p.Arg175His for P3 and P4 hESCs) and c.785G>C (p.Gly262Ala for P3 hESC), were found in the *TP53* DNA-binding domain (DBD), where most *TP53* mutations develop in hPSCs (Amir *et al*., 2017) (Fig. 2A). Remarkably, a missense SNV, c.524G>A, predicted p53 structural damage (Fig. 2A). As previously described (Merkle *et al*., 2017), P4 hESCs with definitive TP53 mutations in DBD become dominant clones soon after p53 stabilization (with Nutlin3a treatment) when mixed with P1 hESCs (Fig. 2B). This result corroborates the observation that p53 protein levels were markedly stabilized in P3 and P4 hESCs even without stimuli (Fig. 2C) with no significant TP53-dependent gene upregulation (i.e., *GADD45A, PPM1D,* and *MDM2*) (Fig. 2D). These findings demonstrate that TP53 mutations in P3 and P4 hESCs were dominant-negative.

**Figure 2.**
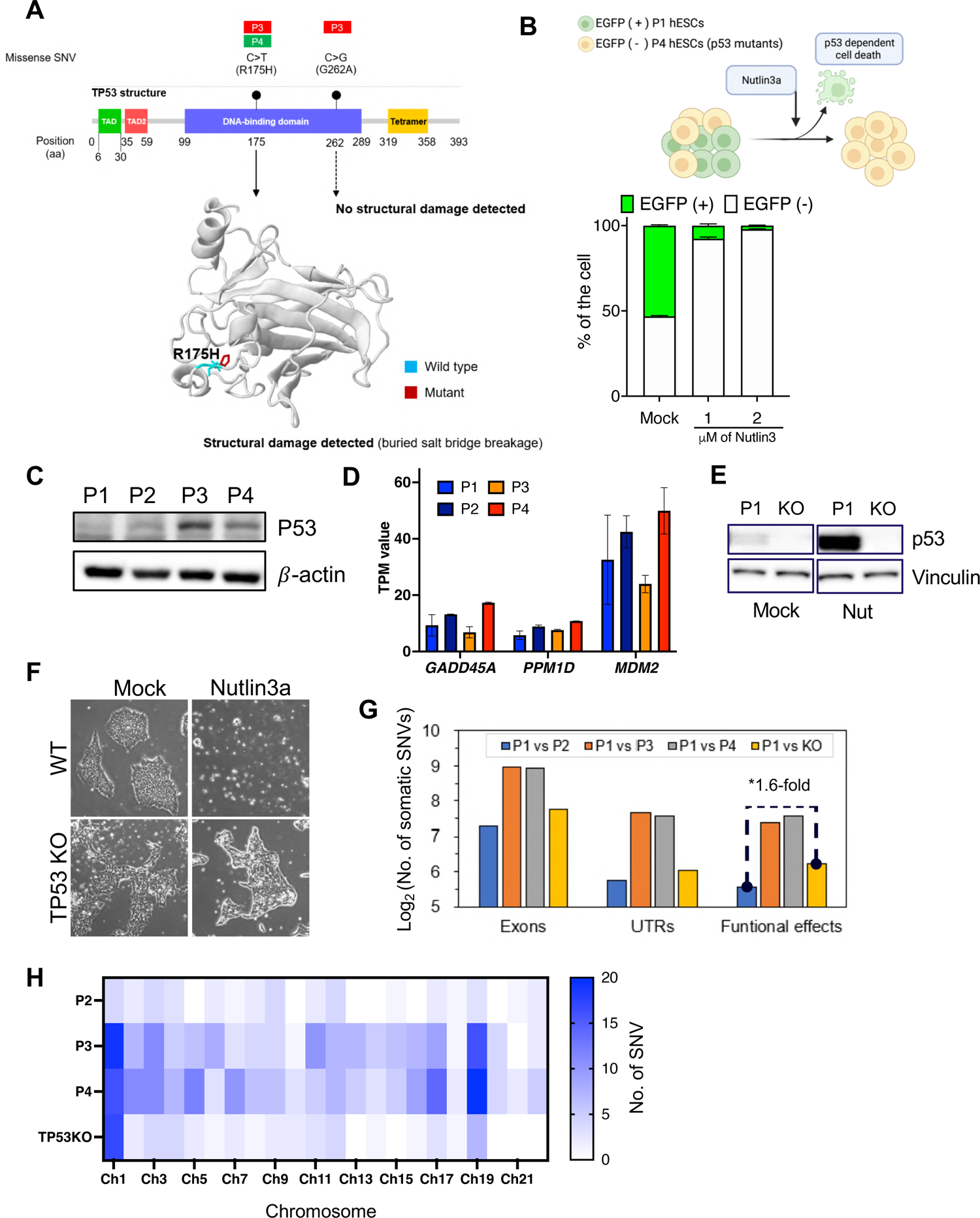
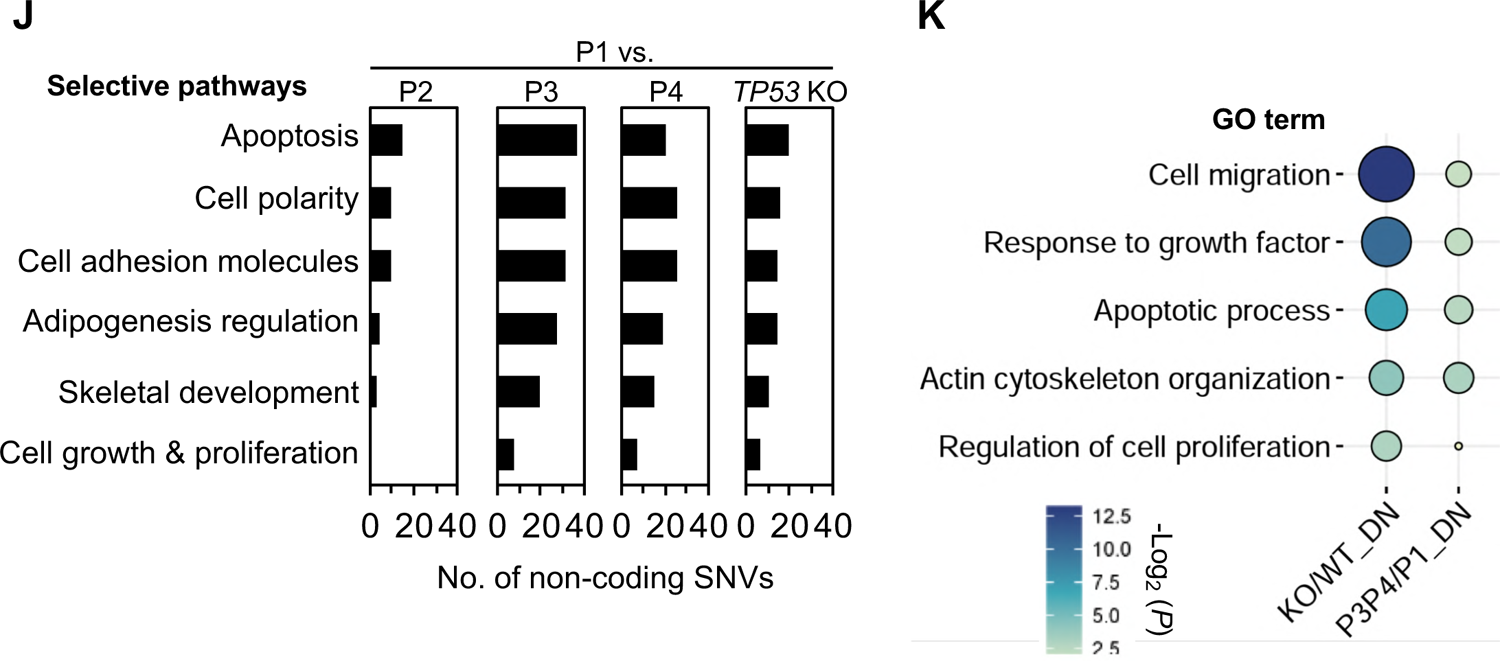
TP53 mutations relative to an increase in somatic mutations (A) The identification of somatic missense SNVs in TP53 of P3 and P4 hESCs and the structural change introduced by the missense SNV (R175H). Structural change of TP53 was predicted using Missense3D. (B) Normal H9 tagged with eGFP and p53 mutant H9 were mixed and cultured. Nutlin3a, the p53 activator, was treated for 24hrs then cell populations were assayed and analyzed by flow cytometry. (C) basal p53 protein level in P1, P2, P3 and P4. (D) Expression levels of p53 downstream genes, *GADD45A*, *PPM1D* and *MDM2*. RNA-Seq for P1, P2, P3, and P4 hESCs was performed, and gene expression levels were quantified by using RSEM with transcripts per million mapped reads (TPM) values. (E) p53 protein level of established TP53KO cell compared to p53 normal H9 were detected after 16hrs of Nutlin3a 1uM treatment by immunoblotting. (F) Resistance against p53 activator Nutlin3a of TP53KO H9 compared to isogenic Mock pair. Nutlin3a 2uM was treated for 24hrs. (G) The number of somatic SNVs identified in exons of P1-derived *TP53* KO hESC. In Figure 2G, untranslated regions (UTRs) encompass both 5’ and 3’ ends, and functional effects include missense, nonsense, and splice sites. (H) The chromosomal distribution of somatic SNVs in P2, P3, P4, and P1-derived *TP53* KO hESCs. The abundance of somatic SNVs was presented with a heatmap. (I) Functional pathway annotation for genes with somatic non-coding SNVs in P2, P3, P4, and P1-derived *TP53* KO hESCs. Functional pathways involved in the functions of survival and proliferation in stem cells related to culture adaptation were selectively analyzed using QIAGEN Ingenuity Pathway Analysis. (J) Gene ontology (GO) enrichment analysis of down-regulated genes in *TP53* KO hESC compared to wild-type (WT or P1). The GO enrichment results were performed and validated as described in method.

In the following STRING network analysis (41), numerous somatic mutations were identified in genes linked to mutated *TP53* in P3 and P4 hESCs (Fig. S2A). This discovery suggests that mutated *TP53* may influence mutations in other gene nodes within the *TP53* network and increase somatic mutations in P3 and P4 hESCs (Fig. 1D). Considering that p53 is integral for genome guidance (Lane, 1992), we theorized that dominant-negative TP53 mutations favor the accumulation of somatic mutations, as depicted in Figure S2A. Therefore, we produced TP53 knockout hESCs (KO hESCs) from P1 hESCs (normal hESCs) by introducing indel (insertion and deletion) in exon 4 with Cas9 (Fig. S2B). The clonal selection after Cas9 established TP53 KO hESCs (KO hESCs) with one base pair (bp) insertion to induce frame-shift mutation (Fig. S2C). The lack of p53 protein (Fig. 2E). Functional p53 KO was verified by the lack of p53 protein and the failure of cell death induction with Nutlin3a in KO hESCs (Fig. 2F).

Through whole-genome sequencing, we identified more exonic somatic mutations in KO hESCs than in P2 hESCs (Fig. 2G). Functional effect variants, including missense, nonsense, and splice-site SNVs (Fig. S2D), were 1.6 times more abundant. The analysis also revealed a predominant distribution of somatic mutations on KO hESC’s chromosome 1, similar to P3 and P4 hESCs (Fig. 2H). This correlation suggests that TP53 mutations accumulate somatic mutations on specific chromosomes. Although most somatic mutations were identified in non-coding KO sequences, we conducted a functional annotation related to cell differentiation for genes with these mutations. The analysis unveiled frequent gene mutations associated with functions such as apoptosis, cell polarity, cell adhesion, adipogenesis regulation, stemness, cytoskeletal development, and chromatin modification, which may be involved when acquiring hESC survival traits (Fig. 2I). In particular, genes related to apoptosis and growth were down-regulated in TP53 KO with loss-of-function (LOF) (Fig. 2J). We also identified numerous somatic mutations in gene nodes within the TP53 KO network (Fig. S2E), indicating somatic mutation expansion and accumulation within the TP53 network. A gain event with abnormal copy number changes all over chromosome 1, with a significant copy number ratio, was also identified (Fig. S2F and S2G). Contrasting P3 and P4 hESCs, CNV was not found at 17q24.1/2 or 20q11.21 in the TP53 KO (Fig. S2H). This result demonstrates that TP53 KO has undergone significant genetic alteration, effectuating genetic instability.

### Culture-adapted hESC cellular heterogeneity and transcriptome profiles

Despite the correlation between LOF TP53 mutations and increased somatic mutations, TP53 mutations could not address the 20q11.21 CNV gain and high representative gene expressions (*TPX2* and *BCL2L1*) to trigger typical cellular events for “culture adaptation” (i.e., abnormal mitosis and survival trait; data not shown) and further genetic aberrations (i.e., additional CNV such 17q24 gain) (Fig. S2H and 3A). Transcriptomes from P1, P2, P3, and P4 hESCs were achieved in single-cell levels to monitor the variations in long-term hESC cultures (Fig. 3B).

**Figure 3.**
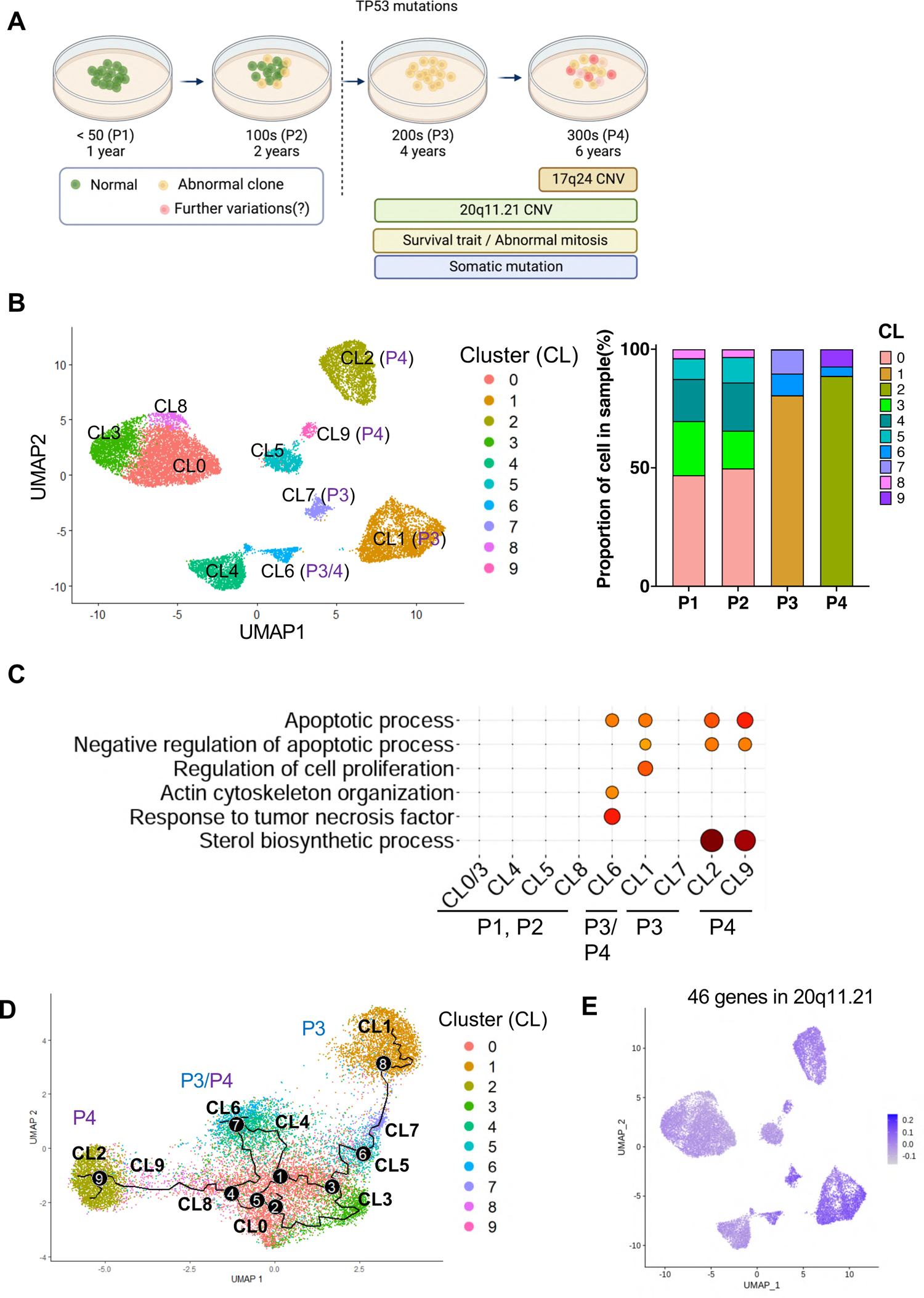
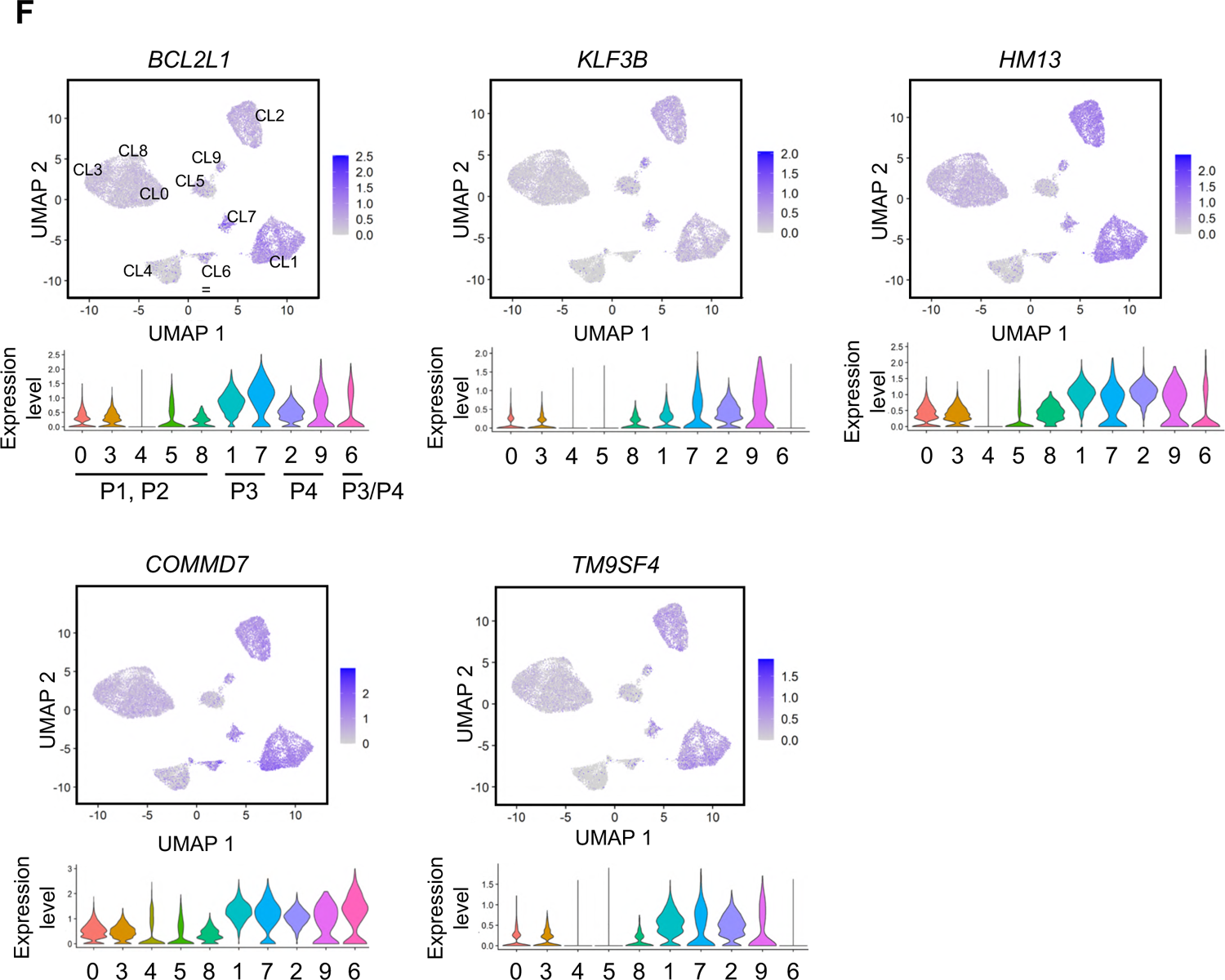
Culture-adapted hESC cellular heterogeneity and transcriptome profiles (A) Scheme of isogenic pair hESCs and single cell analysis considering TP53 mutation status. (B) UMAP plot visualization of cultured hESCs (P1, P2, P3, and P4) colored by 10 different transcriptionally distinct clusters (left panel); CL0, CL3, CL4, CL5 and CL8 for both P1 and P2 hESCs, CL1 and CL7 for P3 hESC, CL2 and CL9 for P4 hESC, CL6 for both P3 and P4 hESCs. The right panel in Figure 3B represents the composition of clusters in each sample and the proportion of cells within each cluster. (C) GO terms enriched for each cluster based on differentially expressed genes. GO enrichment analysis was conducted using DAVID with a cutoff of EASE score < 0.01. (D) Single cell trajectory reconstructed by Monocle 3 for four cultured hESCs. The trajectory indicates that P3 and P4 hESCs have generated significant cellular heterogeneity from P1/P2 hESCs. Even between P3 and P4 hESCs, a high level of cellular divergence is shown. (E) Expression levels of 46 genes in 20q11.21 are indicated in each cell projected on the UMAP plot using the featureplot function. (F) Expression levels of *BCL2L1*, *KLF3B*, *HM13*, *COMMD7*, and *TM9SF4* in 20q11.21 are indicated in each cell projected on the UMAP plot using the featureplot function. The VlnPlot below the UMAP plot shows the distribution of single cell gene expression in each cluster. The y-axis of each panel represents the expression levels of the indicated genes.

The cellular divergence of P3 (clusters 1, 6, and 7) and P4 (clusters 2, 6, and 9) hESCs from P1 and P2 hESCs (clusters 0, 3, 4, 5, and 8) were identified in the UMAP clustering analysis (Fig. 3B). The gene ontology (GO) annotation for genes expressed highly in P3 and P4 hESC clusters revealed a unique function associated with long-term culture adaptation. Gene signatures of the apoptotic process related to negative regulation were consistently altered in P3 and P4 hESC clusters. Similarly, ‘sterol biosynthesis’ was significantly enriched in P4 hESC clusters (Fig. 3C).

The unique enriched geneset in P4 hESCs compared to that of P3 hESCs implied that further variation from P3 hESCs would transpire to acquire the unique P4 hESC characteristics. The RNA velocity analysis to achieve each cluster’s trajectory supported this hypothesis (Fig. 3D). Intriguingly, we noted that genes in 20q11.21 loci, consisting of 46 genes, were relatively increased in all P3 and P4 hESC clusters (Fig. 3E). Expression levels of *BCL2L1, KIF3B, HM13, TM9SF4* and *COMMD7*, which are involved in cell proliferation, differentiation inhibition, and anti-apoptosis (Villa-Diaz *et al*, 2013), in 20q11.21 were remarkably elevated in P3 and P4 hESCs (Fig. 3F), contributing a potent selective advantage in culture.

### Alteration of chromatin accessibility in culture-adapted variants

The survival traits and *BCL2L1* expression evident in variants were not replicated in early-passaged iPSCs carrying the 20q11.21 gain (Jo *et al*., 2020). Transcriptome profiles of iPSCs obtained from the Korea National Institute of Health (KNIH) (eight normal, four with 20q11.21 gain, including four iPSCs previously reported (Jo *et al*., 2020)(Fig. S3A) reinforced this consistency. Notably, the 20q11.21 gain in these iPSCs did not induce gene expressions at the 20q11.21 locus, such as *HM13, ID1, BCL2L1,* and *TPX2* (Fig. S3B). In addition, two early-passaged hESC lines (KR1 and KR2: WA09 hESCs) from two independent laboratories at the Korea Research Institute Bioscience and Biotechnology (KRIBB) were examined. Intriguingly, the KR2 line with a 20q11.21 gain (Fig. S3C) did not exhibit *BCL2L1* induction (Fig. S3D) or cell death resistance from YM155 or Nocodazole (Noc) treatment (Fig. S3E). Furthermore, the KR2 line maintained an intact TP53 status (Fig. S3F).

According to the scRNA data indicating active gene expression at 20q11.21 in later passages (Figs. 3E and 3F), an additional molecular event, such as epigenetic alterations (Andrews *et al*, 2022; Bar & Benvenisty, 2019), could transcriptionally activate genes at this locus during *in vitro* culture. This event may directly induce the abnormal phenotypes of ‘culture-adapted variants’ (i.e., survival trait or abnormal mitosis) (Fig. 4A). Therefore, single-cell ATAC-seq was performed to monitor chromatin accessibility at the single-cell level. Major populations depending on these passages were distinctly clustered with varying chromatic accessibility. However, some minor populations existed in the overall distribution (Fig. 4B). Further classification revealed nine subclusters (Fig. 4C), and hESCs with different passage numbers formed their own major cluster (CL); CL2 in P1, CL3 in P2, CL1 in P3, and CL0 in P4.

**Figure 4.**
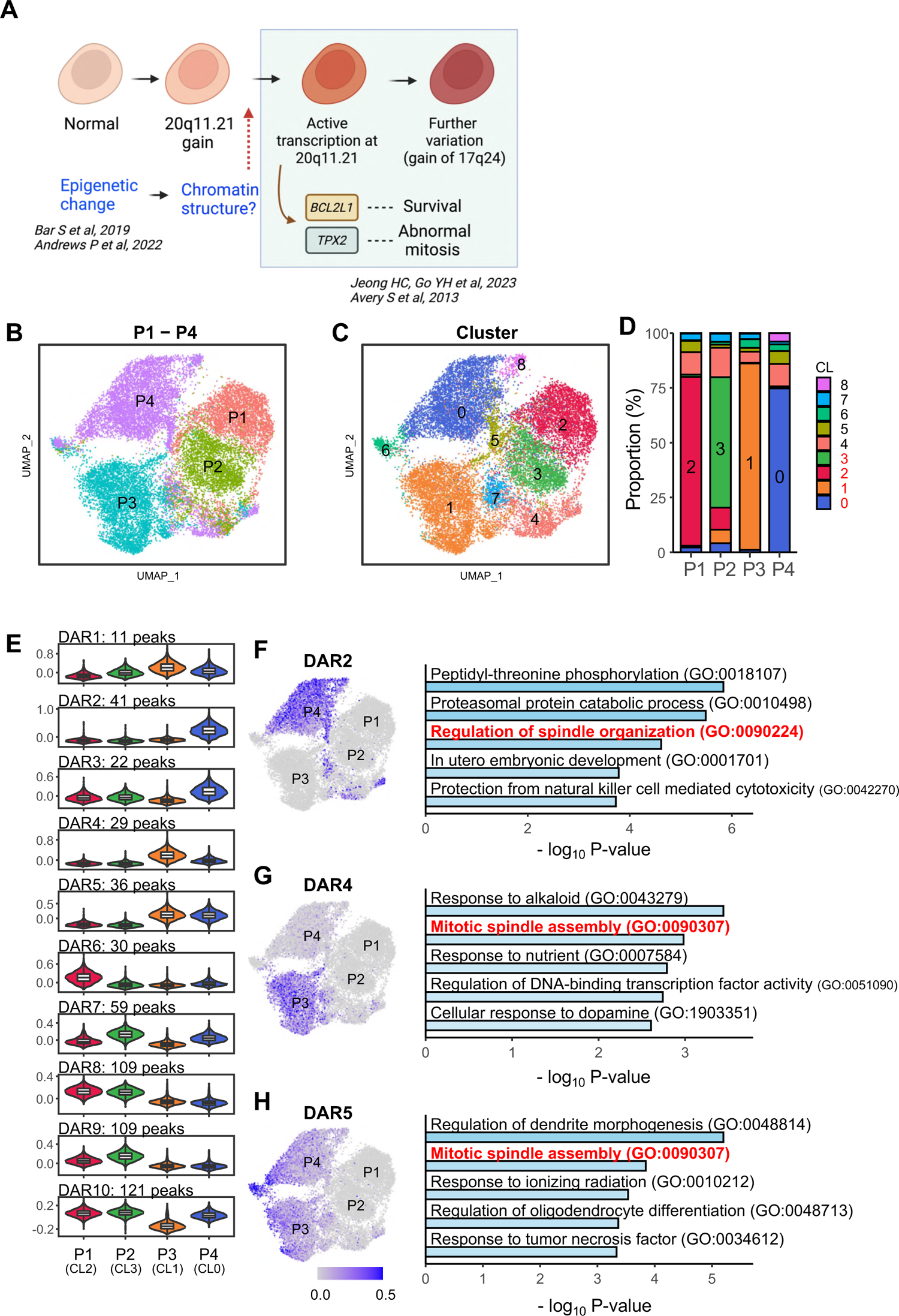
Alteration of chromatin accessibility in culture-adapted variants (A) Scheme of chromatin structural alterations involved in culture-adapted variants. (B) Uniform Manifold Approximation and Projection (UMAP) of scATAC-seq from all samples from P1 to P4. Each dot in the scatter plot represents a single-cell chromatin accessibility. Four samples are color-coded. (C) Clustering analysis depending on the chromatin accessibility. All cells are grouped by 9 clusters and marked by number from 0 to 8. (D) The bar plot represents proportion of a cluster for each sample. The major clusters are CL2 for P1, CL3 for P2, CL1 for P3, and CL0 for P4. These major clusters occupy 80.6% of all cells. (E) The violin plots show average open chromatin levels of differentially accessible regions (DARs) in major clusters. Y-axis represents scaled average open chromatin level of DARs which was calculated by AddModuleScore() function. All DARs were clustered by k-means clustering with k=10. The number of peaks belonging to a specific DAR are indicated. (F-H) Representative DARs corresponding to passages and Gene Ontology analysis using their target genes. The UMAP shows the level of average open chromatin accessibility of indicated DARs. The bar plot represents the top 5 Gene Ontology (GO) terms associated with putative target genes of indicated DAR. The GO terms related with spindle assembly were marked by red. The p-values of the enrichment were calculated by Metascape.

Overall, 567 differentially accessible regions (DARs) were identified in these major clusters. K-means clustering (k=10) grouped all DARs into ten distinct clusters (Fig. 4D). For example, DAR2 and DAR4 chromatin accessibility was predominantly increased in major clusters of culture-adapted variants (i.e., P3 and P4). Thus, potential target genes were selected by the associated cis-coaccessibility network (CCAN) analysis to investigate genes regulated by each DAR and their association with specific phenotypes or cellular processes (Fig. S4A). DAR2, 4, and 5 were generally associated with spindle assembly (Figs. 4E-G), whereas DAR6, 8, and 9 were frequently localized in normal hESCs (i.e., P1 and P2) without common gene ontology terms (Fig. S4B-D). In sharp contrast, DAR2 (primarily in P4), DAR4 (P3), and DAR5 (P3 and P4) were strongly associated with ‘spindle organization’ (Figs. 4E-G).

These results suggest that an epigenetic change that alters chromatin structures regulating genes associated with ‘spindle organization’ is induced in culture-adapted variants (i.e., P3 and P4). Consistently, the aberrant mitosis with a lagging chromosome or chromosome bridge that results from abnormal spindle dynamics occurs in these culture-adapted variants (Jeong *et al*., 2023).

### Epigenetic alteration effects on genetic alteration regarding gene expression

We hypothesized that a molecular event to activate the chromatin structure of 20q11.21 loci would transpire in hPSCs along with a 20q11.21 gain. This event would also promote crucial gene expressions, such as *BCL2L1* (for survival) or *TPX2* (for spindle stabilization and abnormal mitosis), to achieve the typical cellular phenotypes of culture-adapted variants (i.e., survival trait and abnormal mitosis (Avery *et al*., 2013; Jeong *et al*., 2023; Nguyen *et al*., 2014) (Fig. 5A). To verify this theory, subsequent inquiries were conducted to analyze the chromatin structure of loci that underwent increases in DNA copy numbers.

**Figure 5.**
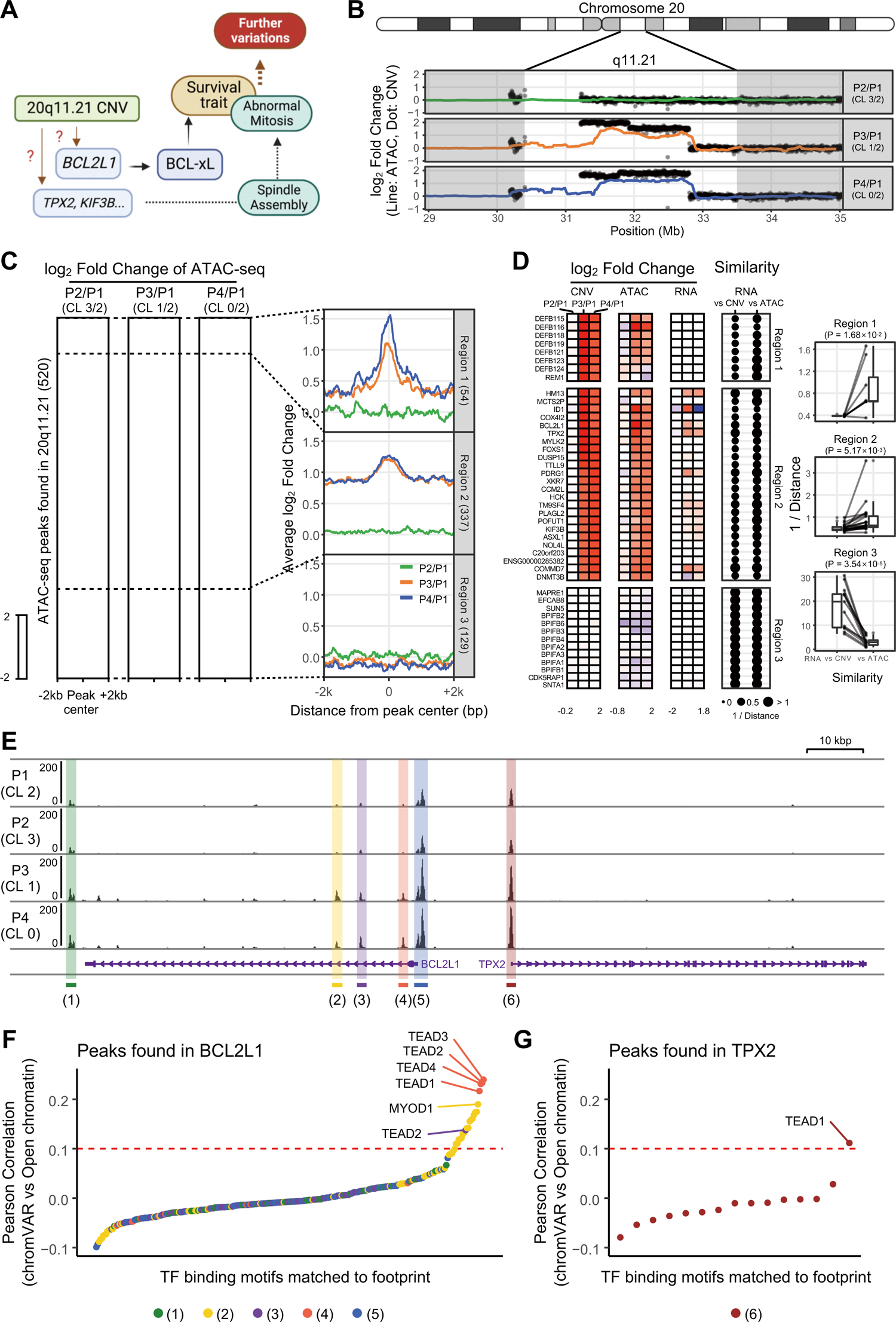
Epigenetic alteration effect on genetic alteration regarding gene expression (A) Scheme of phenotypic variations driven by genetic and epigenetic alterations in culture-adapted cells. (B) ATAC-seq signal along with copy number variation around 20q11.21 locus. The dots and line represent log_2_ fold change against P1 of copy number and ATAC-seq read count for each. (C) The heatmaps represent pattern of log_2_ fold change of ATAC-seq signal around all peaks found at 20q11.21. The fold changes were calculated against P1 major cluster (CL2). The line plots show average log_2_ fold change around peaks found at the same genomic loci. Each region (Region 1, 2, and 3) was separated by dotted lines. (D) Comparative analysis of similarities between CNV and RNA, and between ATAC and RNA. The heatmaps indicate log_2_ fold change of CNV, ATAC and RNA for each gene located at 20q11.21. The dot size represents similarity score calculated by inverse of distance. The significance of similarity score difference was calculated by paired t-test. (E) A representative scATAC-seq profile at BCL2L1 and TPX2 loci. The y-axis value represents normalized ATAC-seq read count. The DAR peaks are marked by color box and numbered. The location and direction of genes were shown at the bottom. (F-G) Pearson correlation of transcription factor (TF) activity score (chromVAR) of transcription factor and open chromatin level at the peak. TF binding motifs were ordered by the correlation value. The TEAD family TF binding motifs show the highest correlation in BCL2L1 locus (Figure 5E (1)∼(5)). TF with the highest correlation in TPX2 locus (Figure 5E (6) is identified as TEAD1.

Open chromatin levels exhibit dynamic changes across the genome, but these changes are dependent on the CNV changes in regions with significant copy number variations (e.g., 20q11.21 in P3 and P4; 17q24.1 and 17q24.2 in P4; Figs. 5B, S5B and S5C). This observation could be attributed to the nature of the ATAC-seq technique, which reads DNA derived from open chromatin regions and enables the measurement of chromatin accessibility.

However, when examining the average open chromatin around peaks at 20q11.21, three distinct patterns were observed (Fig. 5C): Pattern 1 (Region 1, chr20:30,400,000-31,502,846) exhibited a significant increase in open chromatin at the peak center; Pattern 2 (Region 2, chr20:31,502,846-32,804,612) displayed an overall increase in open chromatin around ±2kb of the peak center; Pattern 3 (Region 3, chr20:32,804,612-33,500,000) indicated no changes in CNV and open chromatin.

The broad increase in open chromatin observed around ±2kb of the peak center in Region 2 is attributable to DNA copy number amplification. In contrast, the localized increase in open chromatin at the peak center suggests additional epigenetic alterations. Notably, genes located in Region 2 exhibit increased open chromatin spanning their gene bodies, with a further increase in open chromatin downstream of 1kb from the transcription start site (TSS) (Fig. S5A). This finding implies that gene copy amplification, including regulatory sequences, and additional downstream epigenetic alterations were apparent at the promoter region. Therefore, CNV patterns, open chromatin, and gene expression changes were compared across protein-coding genes within the 20q11.21 locus to investigate the importance of downstream genetic and epigenetic alterations in gene expression (Fig. 5D).

Although there were individual gene-specific differences, patterns in RNA expression changes in Regions 1 and 2 closely resemble the pattern of open chromatin changes than CNV changes. This observation suggests that gene expression alterations in the 20q11.21 locus are likely due to upstream epigenetic alterations. On the other hand, regarding the 17q24 locus, chromatin structure alterations did not resemble RNA expression variations when compared to copy number changes. (Figs. S5D and S5E). This scenario implies that DNA copy number alteration drives chromatin structure changes at the 17q24 locus.

Next, Tn5 footprinting was performed within each cluster (Fig. 5E), and TF binding motifs in the footprints were examined to identify transcription factors (TFs) involved in *BCL2L1* and *TPX2* expression (Figs. 5F and 5G). Among various candidates, the TEAD family exhibited a potent correlation between open chromatin and TF activity. Moreover, the TEAD family’s TF activity consistently ranked higher in P3 and P4 (Fig. S5F). These findings suggest that the TEAD family of TFs, consistent with YAP activation (or the loss of Hippo signaling) in culture-adapted variants (Kim *et al*., 2023; Price *et al*., 2021; Weissbein *et al*., 2019), may be pivotal in achieving culture-adapted phenotypes.

## Discussion

The unique cellular features of hPSCs, characterized by high sensitivity to DNA damage-induced apoptosis and robust DNA damage repair systems, contribute to well-maintained genome integrity (Weissbein *et al*, 2014) and lower mutation rates during *in vitro* culture compared to adult stem cells or organoids (Kuijk *et al*., 2020). Other than chemical supplements (Thompson *et al*., 2020), long-term exposure to high oxygen concentrations during culture has been recently identified as a primary cause of spontaneous mutation (Kuijk *et al*., 2020). Dominant-negative mutations in TP53, in which intact activation drives rapid apoptosis, are rare but recurrent in hPSCs (Merkle *et al*., 2017). In contrast, recurrent copy number variations (CNVs) at specific loci, especially 20q11.21 gain, are relatively common (International Stem Cell *et al*., 2011). The recent meta-analysis of 107 different studies revealed that chromosomes 20, 12, 17, X, and 1 are prevalent hotspots for genetic abnormalities (Assou *et al*, 2020). Despite emerging reports on altered cellular phenotypes, such as acquired survival traits (Avery *et al*., 2013; Cho *et al*., 2018; Nguyen *et al*., 2014), aberrant mitosis (Jeong *et al*., 2023), aneuploidy (Ben-David *et al*., 2014; Na *et al*, 2014), and tumor development (Moon *et al*, 2011; Yamamoto *et al*, 2022), the underlying molecular processes for acquiring “culture-adapted phenotypes” or minimum risk assessments remain poorly understood.

As previously described (Kuijk *et al*., 2020; Thompson *et al*., 2020), somatic hESC mutations were less evident in P2 hESCs (cultured over 100 passages), which exhibit normal copy numbers and the TP53 wild-type. However, somatic mutations, including COSMIC Tier1 mutations, were drastically increased concurrently with TP53 mutations and 20q11.21 CNVs in P3 and P4 hESCs (Fig. 1). The amplification of somatic mutations and gain, especially at chromosome 1 after TP53 knockout (Fig. 2 and S2G), implies that TP53 mutants undergo further genetic abnormality. These results highlight the rationale for avoiding rare TP53 mutant hPSC use in clinical applications.

On the other hand, hPSCs with 20q11.21 gain acquire the survival trait through *BCL2L1* expression. However, *BCL2L1* induction did not manifest in iPSCs or hESCs with definitive 20q11.21 gain during the early passage (Jo *et al*., 2020) (KR2 in Figure S3). Thus, gene expressions at 20q11.21 that induce biological consequences (i.e., survival trait or aberrant mitosis) require additional molecular events. Multi-omics analyses (i.e., scRNA-seq; Fig. 3 and scATAC-seq; Fig. 4) revealed that additional epigenetic events to open the 20q11.21 chromatin structure are necessary for the transcriptional activation of prominent genes at this locus (e.g., *BCL2L1* and *TPX2*) through TEAD upon promoter binding.

Interestingly, the absence of these genetic abnormalities and the similarity of single-cell transcriptome profiles to P1 hESCs in P2 hESCs (despite over 100 passage numbers) suggest that critical events during culture-adapted phenotype development potentially surpass the influence of passage numbers. This finding implies that TP53 mutations or 20q11.21 gain are likely more hazardous than passage numbers. Thus, P3 and P4 hESCs are promising cellular models for monitoring biological consequences after these adverse events regardless of unfeasible passage numbers. Notably, complete TP53 KO increased somatic mutations and induced chromosomal gain, especially on chromosome 1, albeit without concurrent gains at 20q11.21 or 17q.24 (Fig. 2 and S2). Lastly, the current study raises questions regarding the reciprocal nature of these molecular abnormalities, particularly in hESCs (i.e., KR2) with 20q11.21 gain that maintained TP53 wild-type status (Fig. S3F).

In conclusion, our findings emphasize that molecular mechanisms for epigenetic remodeling are crucial for transcriptionally activating genes at the 20q11.21 locus, which induces culture-adapted phenotypes in hPSCs. Based on these results, TP53 mutations and 20q11.21 gain should be regarded as hazardous events rather than passage numbers in hPSCs.

### Data availability

All unique/stable reagents generated in this study will be freely available from the lead contact to academic researchers upon request.

- Whole-genome sequencing (WGS), RNA-Seq, scRNA-Seq data for P1, P2, P3, and P4 hESCs NCBI Sequence Read Archive (SRA) under the BioProject accession number PRJNA1020454(https://dataview.ncbi.nlm.nih.gov/object/PRJNA1020454?reviewer=8qorbhrbs04mrmrmonmm5lek5d).
- scATAC-Seq data for those four samples generated in this study NCBI SRA under the BioProject accession number PRJNA1020991 (https://dataview.ncbi.nlm.nih.gov/object/PRJNA1020991?reviewer=7m7slq0d272jmap4pjsvd2spp9).

## Acknowledgments

This work was supported by a grant from the National Research Foundation of Korea (NRF-2020M3A9E4037904, NRF-2020M3A9E4037905, NRF-2022M3A9B6082674, NRF-2022R1A2C3011663) by Korean Fund for Regenerative Medicine funded by Ministry of Science and ICT, and Ministry of Health and Welfare (Grant number RS-2022-00070316).

## Author contributions

The nature of every author’s contribution was specified using the CRediT contributor taxonomy. H-J. C. T-Y R and C-P.H conceived the overall study design (Conceptualization), led the experiments (Project administration and Funding acquisition), and wrote the manuscript (Writing-original draft, review&editing). Y-J.K. mainly conducted the cellular and biochemical experiments (Data curation), data analysis (Formal analysis) and wrote the specific part of the manuscript (Writing-original draft). S.K and B.K mainly performed WGS, scRNA-seq and scATAC-seq and data analysis (Formal analysis, Data curation), and wrote the specific part of the manuscript (Writing-original draft). S.O, H.L and J-H. J performed sequencing data analysis (Formal analysis, Data curation). D.K established KO lines (Formal analysis, Data curation). D.G and J-H.K performed data analysis of transcriptome and WGS dataset of iPSCs (Formal analysis, Data curation).

## Competing interests

The authors declare that they have no known competing financial interests or personal relationships that could have appeared to influence the work reported in this paper.

## Notes

### Competing Interest Statement

The authors have declared no competing interest.

